# Differential effects of day-night cues and the circadian clock on the barley transcriptome

**DOI:** 10.1101/840322

**Authors:** Lukas M. Müller, Laurent Mombaerts, Artem Pankin, Seth J. Davis, Alex A. R. Webb, Jorge Goncalves, Maria von Korff

## Abstract

The circadian clock is a complex transcriptional network that regulates gene expression in anticipation of the day-night cycle and controls agronomic traits in plants. However, in crops, information on the effects of the internal clock and day-night cues on the transcriptome is limited. We analysed the diel and circadian leaf transcriptomes in the barley cultivar Bowman and derived introgression lines carrying mutations in EARLY FLOWERING 3 (ELF3), LUX1, and EARLY MATURITY 7 (EAM7). Mutations in ELF3 and LUX1 abolished circadian transcriptome oscillations under constant conditions, whereas *eam7* maintained oscillations of ≈30% of the circadian transcriptome. However, day-night cues fully restored transcript oscillations in all three mutants and thus compensated for a disrupted oscillator in the arrhythmic barley clock mutants *elf3* and *lux1*. Nevertheless, *elf3* but not *lux1* affected the phase of the diel oscillating transcriptome and thus the integration of external cues into the clock. Using dynamical modelling, we predicted a structure of the barley circadian oscillator and interactions of its individual components with day-night cues. Our findings provide a valuable resource for exploring the function and output targets of the circadian clock and for further investigations into the diel and circadian control of the barley transcriptome.

## Introduction

The circadian clock is a time-keeping mechanism that reflects the day-night cycle through an endogenous transcriptional rhythm to anticipate dawn and dusk (McClung, 2006). This clock synchronizes internal rhythms with external light and temperature cycles (Harmer, 2009; Greenham and McClung, 2015). The prevalence of circadian rhythms in all domains of life suggests that circadian clocks provide an adaptive advantage for organisms (Edgar *et al*., 2012). The Arabidopsis oscillator contains an interconnected regulatory network of transcriptional repressors and activators (Hsu *et al*., 2013; Fogelmark and Troein, 2014). These components are expressed sequentially to regulate output genes through regulatory elements present in target promoters (Harmer *et al*., 2000; Covington *et al*., 2008; Michael *et al*., 2008b). The Arabidopsis clock is a master regulator of transcription and controls about 30-40% of the global gene expression in a time-of-day specific cycling pattern where transcription of functionally related genes often peaks in clusters (Harmer *et al*., 2000; Covington *et al*., 2008; Michael *et al*., 2008a, Staiger et al. 2013). Expression of such functional clusters often precedes or coincides with the underlying physiological event (Covington *et al*., 2008; Michael *et al*., 2008a), suggesting that circadian control anticipates diurnal regulation to improve physiological performance (Greenham and McClung, 2015).

In Arabidopsis, the circadian system controls many agronomically important processes, such as metabolism, growth, photosynthesis, and flowering time (Greenham and McClung, 2015). Consequently, it has been suggested that the circadian clock is key to improving adaptation and performance of crop plants (Hsu and Harmer, 2013; Bendix et al. 2015). Putative circadian oscillator genes have been identified in the monocot crop barley based on their homology with the Arabidopsis clock genes (Campoli *et al*., 2012; Calixto et al. 2015). Although the circadian oscillator genes diversified via duplication independently between the monocot and eudicot clades, their structure and expression patterns remained highly similar (Campoli *et al*., 2012; Hsu and Harmer, 2013; Bendix et al. 2015). For example, in monocots, the morning expressed MYB-like transcription factor *LATE ELONGATED HYPOCOTYL* (*AtLHY*) CIRCADIAN CLOCK ASSOCIATED 1 (CCA1) is the only ortholog of the Arabidopsis paralogs CIRCADIAN CLOCK ASSOCIATED 1 (CCA1) and *LHY* (Takata *et al*., 2009; Campoli *et al*., 2012). *HvLHY* overexpression in Arabidopsis causes arrhythmia, suggesting circadian functionality (Kusakina *et al*., 2015). *CCA1* and *LHY* suppress the *PSEUDO RESPONSE REGULATORs* (*PRRs*), which duplicated independently from three ancient *PRR* genes after the divergence of monocots and eudicots such that the orthologous relationship within the *PRR3/7* and *PRR5/9* clades of Arabidopsis and monocot plants cannot be immediately resolved (Takata *et al*., 2010). Partial complementation of Arabidopsis *prr7-11* by *HvPRR37* suggests that the barley gene might retain some functionality of the Arabidopsis orthologue (Kusakina *et al*., 2015). However, *PRR37* orthologs in monocots, *PPD1* in barley and wheat (Turner *et al*., 2005; Beales *et al*., 2007) and *SbPRR37* in sorghum *(Sorghum bicolor)* (Murphy *et al.*, 2011), are major determinants of photoperiod sensitivity and flowering time, whereas natural variation in *PRR* genes in Arabidopsis did not have any notable effect on flowering time (Ehrenreich *et al*., 2009). Furthermore, the genes underlying the two *early maturity* mutants, *early maturity 8* (*eam8*) and *eam10*, have been identified as barley homologs of the Arabidopsis clock genes *EARLY FLOWERING 3* (*ELF3*) and *LUX ARRHYTHMO* (*LUX1*), respectively (Faure *et al*., 2012; Zakhrabekova *et al*., 2012; Campoli *et al*., 2013). Mutations in both genes cause photoperiod insensitivity and early flowering under long and short day conditions in barley (Faure *et al*., 2012; Zakhrabekova *et al*., 2012; Campoli *et al*., 2013). In Arabidopsis, these genes form together with EARLY FLOWERING 4 (ELF4) the evening complex that controls rhythmicity of the clock and growth and development. In barley, several *ELF4*-like homologs exist, including *HvELF4-like 4* that can complement an Arabidopsis *Atelf4* null mutant (Hicks et al. 2001, Kolmos *et al*., 2009). While a number of putative clock components in barley have been identified, there is little information on the contribution of the clock versus day-night cues on the global transcriptome in barley.

We generated diel and circadian RNAseq datasets of four barley genotypes, the spring barley Bowman (BW) and three derived introgression lines with mutations in *HvELF3* (BW290), *HvLUX1* (BW284), and *EARLY MATURITY 7 (EAM7*) (BW287) (Faure *et al*., 2012; Campoli *et al*., 2013). The candidate gene for *EAM7* has not yet been identified, but loss of *EAM7* function accelerates flowering by abolishing sensitivity to the photoperiod (Gallagher et al., 1991). We used the RNAseq time-course data to analyse the effects of barley clock genes on diel and circadian transcriptome oscillations including changes in phase and period under constant conditions and light and dark cycles. Dynamical modelling allowed us to predict a molecular structure of the barley circadian oscillator and to uncover how circadian oscillator components interact with day/night cues to regulate the global transcriptome in barley.

## Results

### Diel and circadian oscillations of the barley transcriptome

We analysed the diel and circadian global leaf transcriptome of the barley cultivar Bowman and the derived introgression lines carrying mutations in *HvELF3* (BW290), Hv*LUX1* (BW284) and Hv*EAM7* (BW287). Plants were grown under cycles of 12h light and 12 h night (LD) and the second leaf of replicate plants was harvested every four hours over 24h. Additional samples were taken in a 2h interval at dusk in all genotypes and additionally at dawn in Bowman (Supplemental Figure S1). Thereafter plants were transferred to constant light and temperature conditions (LL) and leaf samples were taken every four hours for 36h starting from the first subjective night. Individual libraries were single-end sequenced on a HiSeq 2500 with 10 Million reads per library and reads were mapped against a custom reference sequence consisting of 68,739 transcripts (Digel et al., 2015). The nomenclature of the gene models used in this study (Digel et al., 2015) were cross-referenced with the identifiers of the HORVU gene models annotated on the barley pseudochromosomes (Mascher et al., 2017). Raw read counts normalized to counts per million (CPM) were used for the downstream rhythmic analysis and modeling. We determined the oscillating patterns of gene expression including period, as the duration of one complete oscillation and phase as the timepoint of transcript peak expression (Wu et al., 2016; Yang et al., 2010). To increase the analytical power for the rhythmic analysis, the 24 h diel dataset, but not the 36 h LL data, was duplicated to imitate 48 h of sampling data. We identified 18,500 transcripts with expression levels greater than 5 counts per million (cpm) in at least two libraries. Among 18,500 transcripts expressed across all the investigated lines, 84% were scored rhythmic under LD in Bowman (Figure 1, Supplemental Data 1). Under LL, about 23% of the 18,500 transcripts were rhythmic, which is a distinctive feature of clock-regulated genes (Figure 1, Supplemental Data 1). The gene ontology (GO) analyses revealed that, in Bowman under LL, the circadian-controlled transcripts were primarily related to the processes of regulation of DNA-dependent transcription, translation, electron transport, signal transduction, responses to salt stress and cold, and metabolic processes, including amino-acid, sucrose and starch metabolism (Figure 2e, Supplemental Data 1). The molecular functions of the circadian controlled transcripts in Bowman in LL were primarily represented by protein, zinc ion and ATP binding, DNA and nucleotide binding, and sequence-specific DNA-binding transcription factor activity GO terms (Figure 2e).

**Figure 1:**
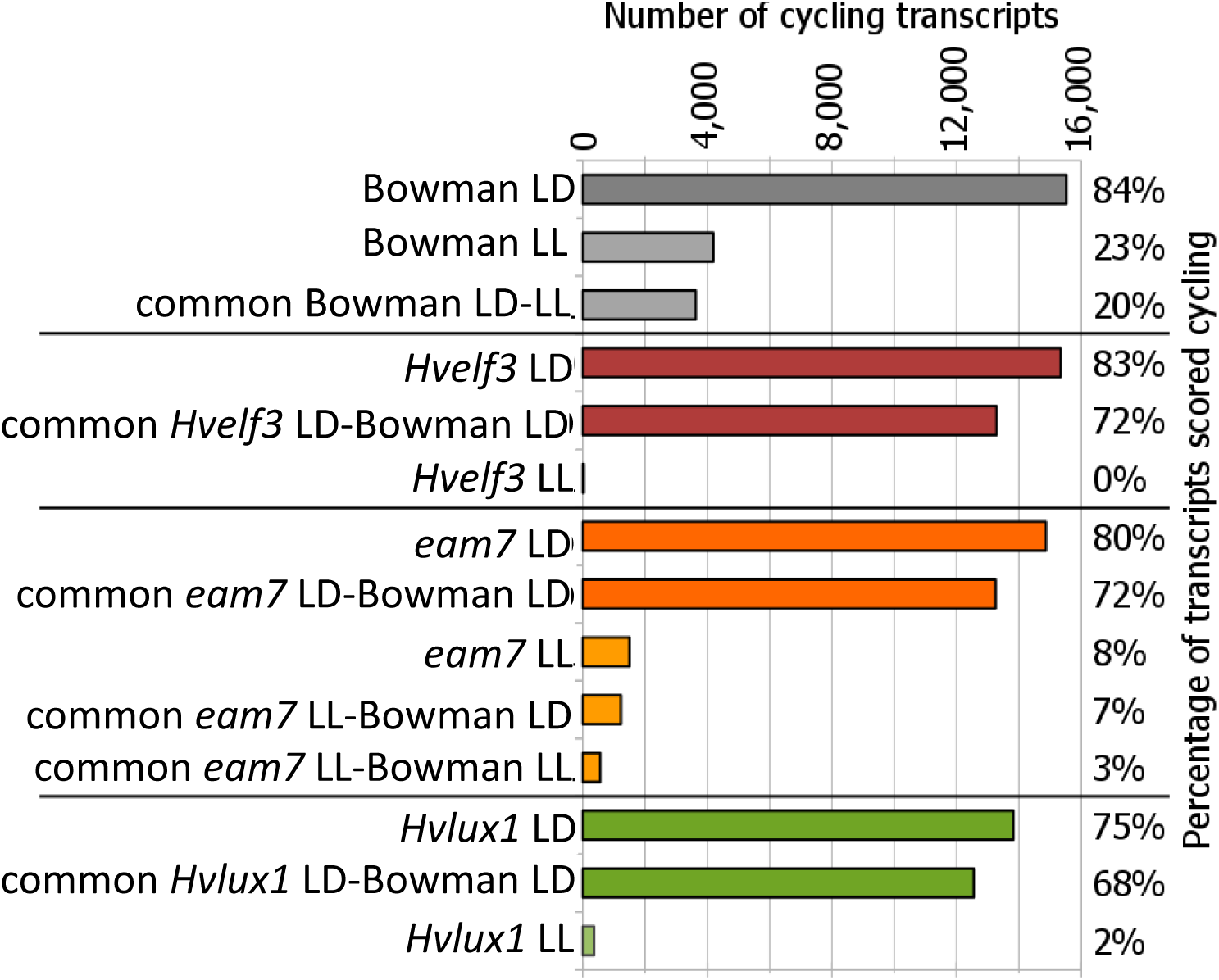
Fraction of transcripts with oscillating transcription pattern. BW WT: Background Bowman; BW290 (*Hvelf3*), BW287 (eam7) and BW284 (*Hvlux1*): Clock mutant genotypes in the Bowman background; ND: night/day cycles; LL: free-running conditions of constant light and temperature. Fractions refer to a total of 18,500 transcripts expressed in all genotypes.

**Figure 2:**
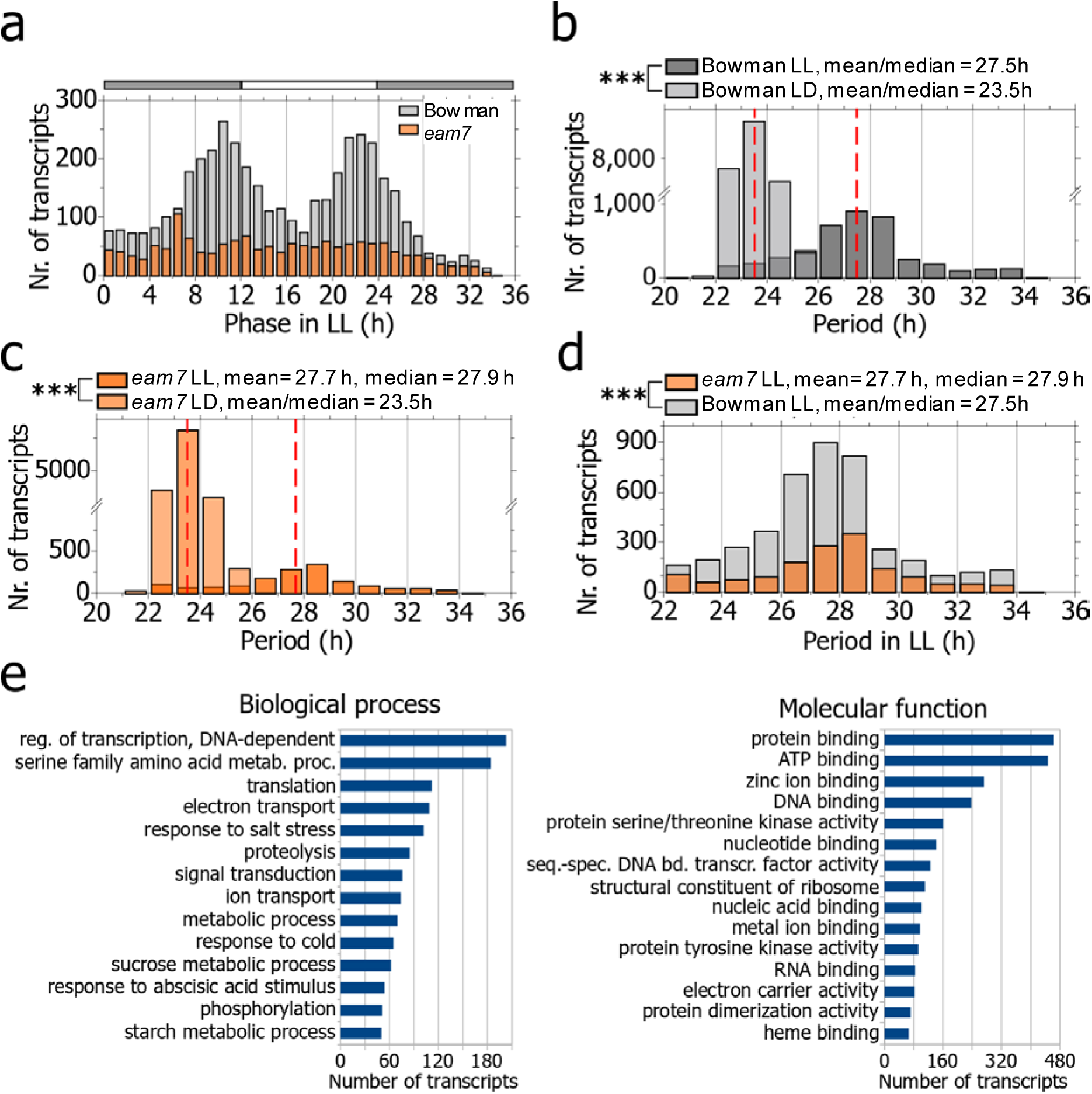
Distribution of the period and the phase of the oscillating transcriptome in constant light and their involvement in biological processes and molecular functions. Phase distribution in constant light (LL) of Bowman (BW WT) and BW287. Grey-white bars indicate the subjective night (grey) and subjective day (white) in constant light conditions. **b, c)** Period distribution of the oscillating transcriptome under constant light (LL) in comparison with night/day cycles (ND) in b) Bowman wild-type (BW WT) and c) BW287. **d)** Comparison of the period distribution in constant light (LL) between Bowman (BW WT) and BW287. **e)** Top-15 categories of the GeneOntology terms for biological processes and molecular function of the transcripts oscillating in Bowman (BW WT) under constant light (LL).

We found that the majority of the transcripts expressed rhythmically under LL were also rhythmic under LD (20% of all the transcripts, 87% of LL transcripts). This demonstrated that about one-quarter of the Bowman transcriptome is modulated by the circadian clock, whereas the largest proportion of the rhythmic transcripts in LD required day-night cues for their rhythmic expression.

The large impact of external transitions on transcriptome oscillations independent of the clock was further supported by the analysis of the *Hvelf3* plants. In *Hvelf3*, no transcript rhythms were detected under LL demonstrating that a functional *HvELF3* is required for self-sustained transcriptome oscillations in barley (Figure 1, Supplemental Data 1). Environmental cues under LD restored oscillatory dynamics in the *Hvelf3* loss-of-function line with 83% of the global transcriptome being rhythmic in the *Hvelf3* plants (Figure 1). The number and the identity of oscillating transcripts were comparable between *Hvelf3* and Bowman plants under diel cycles (Figure 1). In *Hvlux1* plants, only 2% of the expressed transcripts were rhythmic under LL suggesting that, like Hv*ELF3,* Hv*LUX1* is required for free-running oscillations under LL (Figure 1). Once again, LD cycles were sufficient to restore transcriptional rhythms in the Hv*lux1* mutant, i.e. 75% of the transcriptome oscillated in *Hvlux1* plants under LD (Figure 1). Mutation of the *EAM7* locus in BW287 reduced the pervasiveness of circadian transcriptional oscillations, but did not completely abolish them because 8% of the expressed transcripts cycled under LL in *eam7*, about a third of the number of the oscillating transcripts in Bowman (Figure 1, Supplemental Data 1). Under LD, 80% of the global transcriptome was rhythmic in *eam7* and 72% of the rhythmic transcripts were common between *eam7* and the Bowman plants (Figure 1).

Our data demonstrate that cycles of light and temperature and the circadian oscillator drive rhythmic expression in barley. Hv*ELF3, HvLUX1* and *EAM7* contribute to free-running oscillations under constant conditions while environmental rhythms are sufficient to drive rhythmic expression in the absence of a free-running oscillator.

### *EAM7* is a modulator of a bimodal phase distribution under LL and shortens the free-running period

To investigate temporal expression patterns of the circadian-regulated transcripts under free-running conditions, we estimated the phase and the period of every circadian-regulated transcript in the two genotypes that sustained free-running circadian rhythms, Bowman and *eam7*. In Bowman, the distribution of the circadian transcriptome expression phase followed a bimodal pattern with the highest number of transcripts peaking shortly before the transitions to subjective days and nights (Figure 2a, Supplemental Data 1). By contrast, in *eam7* this phase pattern of the cumulative circadian transcriptome was not evident (Figure 2a). These findings indicated that *EAM7* is required to modulate the characteristic bimodal pattern of the circadian transcriptome expression in barley. The period estimates of the oscillating transcripts under LL ranged between 22 h and 34 h in Bowman and *eam7* and followed a bell-shaped distribution with mean periods of 27.5 h and 27.7 h in Bowman and *eam7*, respectively (Figure 2b, c). In both Bowman and *eam7*, the standard deviation of the period distribution was higher under LL (6 h) than under LD (2.5 h) (Figure 2b, c). This could arise from either the uncoupled nature of cellular oscillations in free-running conditions or is a consequence from the period estimation as the signal amplitude was lower in LL than in LD. A longer mean period of oscillating expression patterns in *eam7* suggested that the free-running period under LL was extended in *eam7* compared with Bowman.

### Regulation of the transcriptome-wide phase in day/night cycles

Next, we investigated the transcriptome oscillations under the diel LD conditions. In all genotypes, including those that were arrhythmic in LL, the mean of the period distribution was consistent with the enforced 24-h diel cycle and ranged between 23.5 and 23.6 h (Supplemental Figure S2). The phase was bimodally distributed over the day/night cycle in Bowman so that for the highest number of transcripts the peak of expression occurred before dawn and dusk and, the number of transcripts with the peak expression during the night and day was the lowest (Figure 3a). This pattern was comparable with the phase distribution under LL (Figure 2a) and the transcripts that oscillated in both LL and LD were also bimodally distributed under the diel cycles (Figure 3a). This suggested that the bimodal distribution of transcriptome-wide gene expression is, at least partly, under control of the circadian clock.

**Figure 3:**
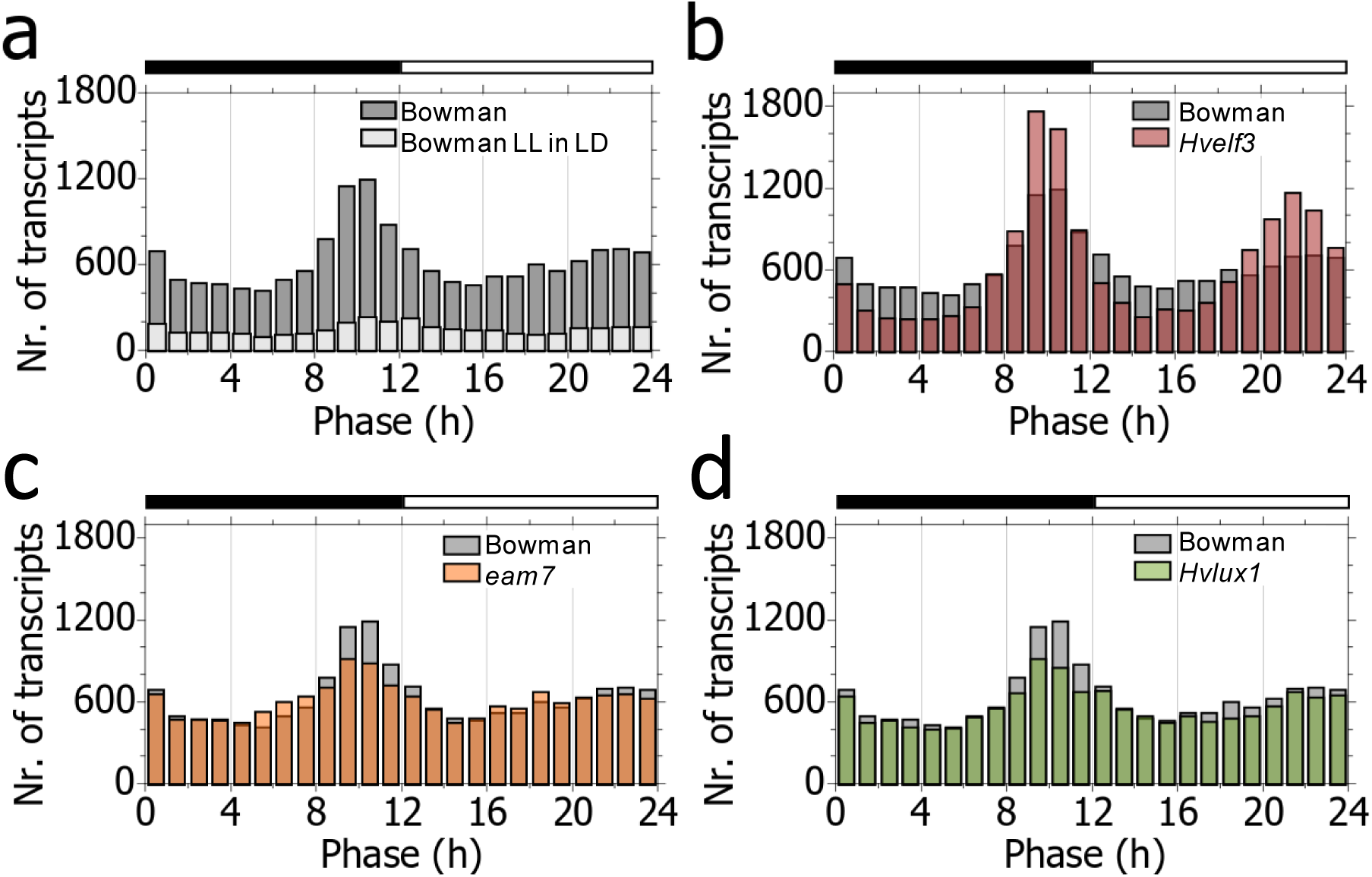
Distribution of the phase of the oscillating transcriptome in night/day cycles. **a)** Phase distribution in Bowman in night/day (ND) cycles for the global oscillating transcriptome (BW WT) and those transcripts detected oscillating in both night/day cycles and constant light (BW WT LL in ND). **b), e), f)** Phase distribution in diel cycles in a) BW290, e) BW287 and f) BW284 in comparison with the Bowman wild-type background (BW WT). **c), d)** Detailed views of Figure 3b. Time point of peak expression (phase) of transcripts that peaked in Bowman (BW WT) but not in BW290 before the c) night-to-day transition (time points between 8h-12h in Figure 3b) and d) the day-to-night transition (time points between 20h-24h in Figure 3b).

The analysis of the clock mutants, however, suggested that the bimodal phase distribution under LD is controlled by both the circadian clock and day/night cues. In *Hvelf3* the phase was bimodally distributed under diel cycles similar to Bowman, however the quantitative characteristics of the phase distribution differed. Namely, in *Hvelf3*, the phase distribution showed higher peaks at dawn and dusk and deeper troughs during the night or the day than in Bowman (Figure 3b). A large number of the transcripts that peaked around the night-to-day and day-to-night transition in *Hvelf3* (Figure 3b) peaked during the day or the night in Bowman. This demonstrated that Hv*ELF3* modulates timing of peak expression of multiple transcripts in day/night cycles. This effect was apparently completely or partially independent of the oscillator defect that causes arrhythmia in the *Hvelf3* plants under LL since the phase distribution in Hv*lux1* mutants under LD was comparable to Bowman (Figure 3c, d), even though self-sustained circadian oscillations were also absent in this genotype under LL conditions (Figure 1). This was also evident from the transcriptome-wide comparison of the phase between the barley clock mutants with Bowman under LD (Supplemental Figure S3). Here, the phase distributions strongly correlated between *Hvlux1* and Bowman (Pearson correlation ρ=0.97, R^2^=0.94) while the phase distributions in *hvelf3* and Bowman were correlated to a lower degree (Pearson correlation ρ=0.93, R^2^=0.86), even though both mutant genotypes harbor an arrested oscillator under LL conditions (Figure 1).

Day/night cycles had strong effects on the phase distribution of the transcriptome as demonstrated by the analysis of the *eam7* transcriptome. Whereas the phase distribution was not bimodal in *eam7* under LL (Figure 2a), under LD, the phase distribution was bimodal similar to the one in Bowman (Figure 3c). Consistently, the phase distributions under LD were highly correlated between *eam7* and Bowman (Pearson correlation ρ=0.96, R^2^=0.92, Supplemental Figure S3). Consequently, external cues under LD controlled the phase of the global transcriptome in *eam7* to peak at the night/day transitions despite the circadian defects observed in *eam7* under LL. Together, these results demonstrated that the bimodal distribution of the phase in diel cycles is controlled by both day/night cues and the clock component Hv*ELF3*. The genetic defects and their underlying circadian phenotypes in *hvlux1* and *eam7* have limited effects on the phase of the global oscillating transcriptome in diel cycles despite their strong transcriptional phenotypes under LL.

### Dynamical models predict components and regulatory interactions of the barley clock

We then sought to infer the regulatory relationships between components of the barley circadian clock. To this end, we modeled a transcriptional network based on the RNAseq time-series data. Our data suggested that Hv*ELF3* and Hv*LUX1* are integral components of the barley oscillator as they were necessary to sustain transcriptome oscillations under LL (Figure 1). Therefore, we hypothesized that modeling a transcriptional network around Hv*ELF3* and Hv*LUX1* could identify the regulatory relationships that shape the circadian clock in barley. We followed an approach that searches the dynamic dependencies of Hv*ELF3* and Hv*LUX1* expression on other transcripts. We used Linear Time Invariant (LTI) models, for interpreting expression data without relying on *a priori* knowledge of the transcriptional network (Dalchau *et al*., 2010; Herrero *et al*., 2012; Mombaerts et *al.*, 2019; Supplemental Information). LTI models require transcriptional data sets that display robust changes in expression over time under free-running conditions. Therefore, only the expression datasets from Bowman and *eam7* could be used in modeling, since their transcriptomes oscillated under LL conditions. In both Bowman and *eam7*, the transcripts encoding Hv*ELF3* had a very low signal-to-noise ratio due to low rhythmicity under LL and could not be used for modeling. We therefore rooted the network around *HvLUX1*, which displayed robust oscillatory dynamics (Supplemental Figure S4). To reduce the identification of erroneous interactions we filtered all circadian transcripts for those that were homologous to Arabidopsis genes representing transcription factors that were labeled “circadian”, thus show circadian expression but are not necessarily components of the core clock (www.geneontology.org). Indeed, while our modeling methodology is computationally inexpensive, the uncertainty about the structure of the network is increasing exponentially with the number of genes considered. Additionally, we filtered the resulting 131 transcripts (Supplemental Data 2) for those that exhibited unambiguous dynamics and a high signal-to-noise ratio of expression in both Bowman and *eam7*. This filter was applied because of the transitional nature of constant light data, which typically shows a large decrease of amplitude after few hours in barley, and the dependency of noise on gene expression levels. *Hvlux1* and *Hvelf3* datasets were not considered in the following network analysis since these mutations led to the arrhythmic transcriptomes. This resulted in 42 transcripts in Bowman and 41 in *eam7* of which 35 transcripts were in common between the Bowman and *eam7* and used for modeling (Supplemental Data 3). For the 35 shared gene transcripts we predicted all possible pair-wise (or single source-target) dynamic dependencies based on the transcript abundance over time for both the Bowman and the *eam7* datasets. For each of these 35 pair-wise comparisons, we fitted a model that captures the expression pattern of Hv*LUX1* in each of the two genotypes.

We then investigated the consistency between the models obtained for Bowman and *eam7* using the v-gap metric (Supplemental Data 4, Supplemental Figure S5). This approach estimates differences between models and allowed us to identify regulatory interactions that were maintained or abolished in the *eam7* mutant (Mombaerts *et al*., 2019). Following this approach, we identified 20 transcripts and 79 regulatory links in Bowman of which 15 transcripts and 49 regulatory links could be cross-validated in *eam7* (Figure 4, Supplemental Figure S5, S6, Supplemental Data 4). Five transcripts could not be confirmed in *eam7* either because they were arrhythmic in *eam7*, the regulatory link could not be modelled with high confidence (model fitness) or the models for Bowman and *eam7* displayed a large difference (v-gap). Among the five genes that could not be confirmed in *eam7* transcripts with homology to REVEILLE 8/6/4 (RVE8/6/4, Hv.12868, HORVU7Hr1G001830.3) and RVE1 (Hv.25709, HORVU6Hr1G066450.5) were the most prominent with eight and seven links to putative clock genes in Bowman, respectively. Both transcripts did only display rhythmic oscillations under LL in Bowman, but were arrhythmic in *eam7*. Among the 15 cross-validated candidates, nine transcripts were encoded by barley genes homologous to known Arabidopsis core oscillator genes. In addition to HvLUX1 as a core of the model (Hv.20312, HORVU3Hr1G114970), the predicted components of barley circadian clock were barley homologs of *LHY* (Hv.5253, HORVU7Hr1G070870) (Mizoguchi *et al*., 2002), REVEILLE 8 *(RVE8)* (Hv.6145, HORVU6Hr1G066000) (Hsu *et al*., 2013), *PRR95* (Hv.4918, HORVU5Hr1G081620) (Farre *et al*., 2005), *PRR59* (Hv.18813, HORVU4Hr1G021000) (Nakamichi *et al*., 2010), *FLAVIN-BINDING, KELCH REPEAT, F-BOX 1* (FKF1) (Hv.4076, HORVU7Hr1G099010) (Baudry *et al*., 2010), GIGANTEA (*GI)* (Hv.1530, HORVU3Hr1G021140) (Dalchau *et al*., 2011), and ZEITLUPE (*ZTL*) (Hv.10907, HORVU6Hr1G022330) (Más *et al*., 2003). Such a result supports that circadian-clock genes are themselves strongly driven by circadian processes.

**Figure 4:**
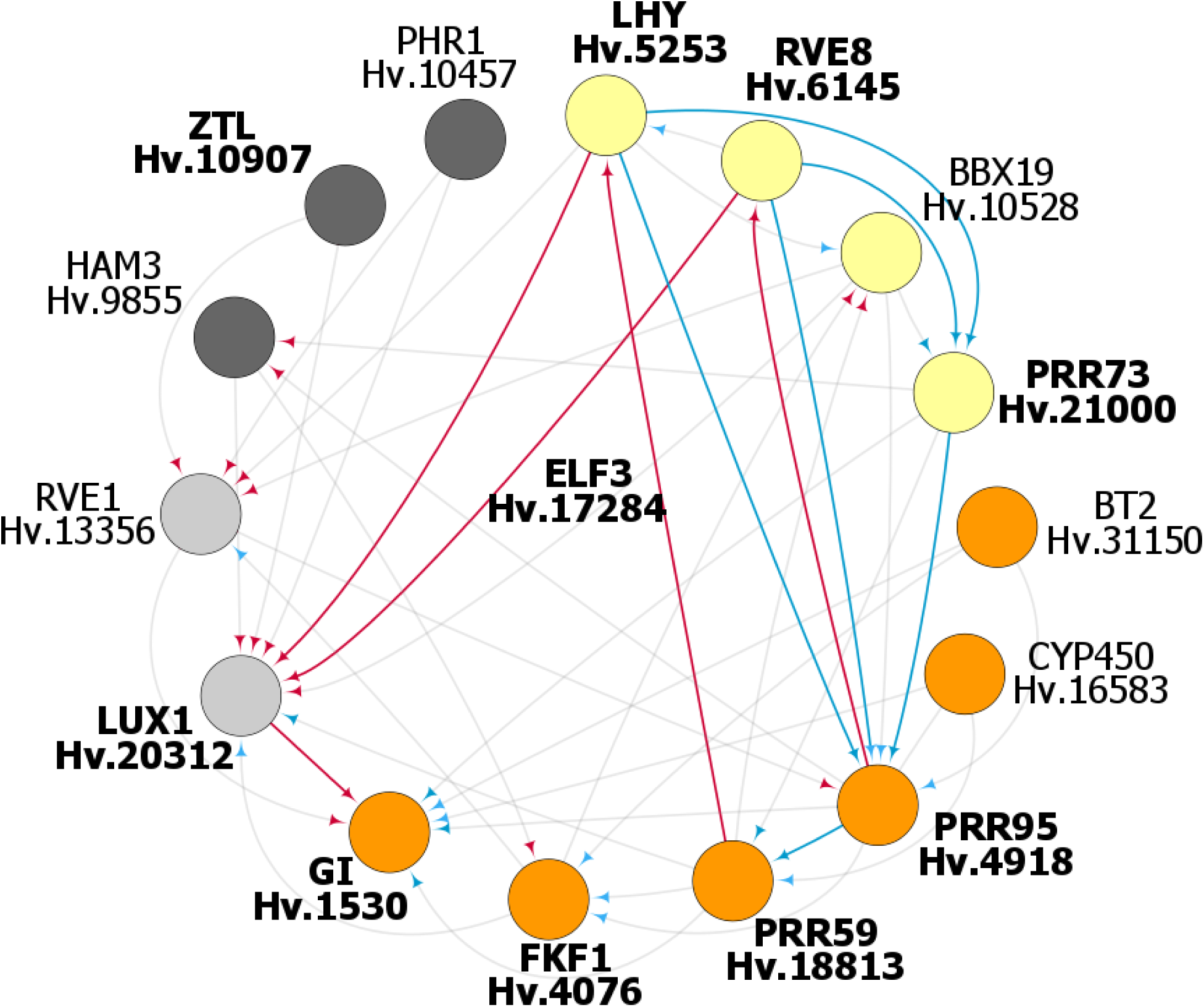
The putative circadian network of the barley oscillator as predicted from time-series expression data. Genetic evidence but no model prediction allowed placing HvELF3 as a core clock component. The figure displays the inferred components and interactions that constitute the barley circadian transcriptional network (also see Supplemental Information). Circadian clock components are represented by circles and sorted in clockwise direction for the time point of peak expression starting with HvLHY at dawn (yellow: morning, orange: evening, grey: night). The regulatory interactions are represented by directed arrows, where activation is marked in blue and inhibition in red. The components printed in bold and the links highlighted in color are consistent with key components and key regulatory principles present in circadian clock models from Arabidopsis.

On the resulting conjunction network, we noted that HvPRR95 appeared as a hub with eight connections, whereas HvLUX1 as the origin of the graph had eleven connections. This indicated a significant role of HvPRR95 in the regulation of the core circadian genes. Therefore, we repeated the search for regulators of HvPRR95, computed their interactions in both datasets, and tested for their consistency. Consequently, four genes were added to the final network REVEILLE-like 7 (HvRVE7, Hv.13356, HORVU2Hr1G104580), Cytochrome P450 (Hv.16583), HvPRR73 (HORVU4Hr1G057550) (Farre *et al*., 2005), and BTB/POZ and TAZ domain-containing protein 2 (HvBT2, Hv.31150, HORVU3Hr1G092090) (Figure 4; Supplemental Data 5). Our modeling did not place barley homologs of other known Arabidopsis clock genes *TOC1*, *ELF4*, Hv*ELF3,* and *PRR37* in the barley clock model. In the case of Hv*TOC1* and barley homologs of *ELF4*, the inferred regulatory interactions to other putative clock components were weak and did not pass the cutoff-filter. The weak transcript oscillations of *HvELF3* precluded its modeling as part of the barley clock. However, we placed HvELF3 in the core clock model based on the genetic evidence that HvELF3 is required for clock function in barley (arrhythmic phenotype of *Hvelf3* mutant). Furthermore, *PRR37*, the photoperiod response gene *Ppd-H1*, did not display rhythmic oscillations under LL and was therefore not placed into the circadian oscillator. Based on the timing of the peak expression starting with *HvLHY* expression at subjective dawn, we arranged the predicted components into a model of the barley circadian clockwork (Figure 4).

In addition to the barley homologs of known Arabidopsis oscillator genes, our analysis suggested several previously uncharacterized components of barley circadian clock. These included the B-Box Zinc Finger Protein 19 (Hv*BBX19*, Hv.10528, HORVU5Hr1G081190) and Hv*RVE7* (Hv.13356, HORVU2Hr1G104580) (Figure 4). In our model, both Hv*BBX19* and *HvRVE7* regulate Hv*PRR95* and are regulated by Hv*LHY* (Figure 4). The modeling predicted that Hv*RVE7* represses Hv*PRR95* and Hv*BBX19* activates Hv*PRR73* and Hv*PRR95*. Other predicted components of barley circadian clock were a homolog of *HAIRY MERISTEM3* (*HAM3*) (Hv.9855, HORVU6Hr1G063650), of BT2, *CYTOCHROME 450* (*CYP450*) (Hv.16583, HORVU2Hr1G025160), and *PHOSPHATE STARVATION RESPONSE 1* (*PHR1*) (Hv.10457, HORVU4Hr1G051080). However, all of these genes were predicted to regulate clock components, but were not themselves regulated by the clock genes (Figure 4). To summarize, our analysis was able to predict components of the barley clock, which are close homologs of the Arabidopsis clock genes (Campoli *et al*., 2012; Calixto et al. 2015) and additionally identified HvBBX19, HvRVE7 and HvHAM3 as putative components of the barley clock.

### Relationship between internal and external cues to regulate the global transcriptome in barley

To quantify the relationship between the circadian oscillator and light signaling in regulating the rhythmicity of barley transcripts, LTI models that integrate both inputs explicitly were computed for each transcript. As a morning clock gene, the expression pattern of *HvLHY* accounted for the contribution of the circadian oscillator, while the light/dark cycle was integrated as a rectangular input (1 = light ON, 0 = light OFF) (Supplemental Figure S7). The expression pattern of the output transcript was approximated by finding the combination of inputs that fit the data best. Then, the contribution of each input was formally compared using a Bode analysis (Dalchau *et al*., 2010). The analysis estimated that 43% of the transcripts that oscillate in both day/night cycles and constant light were predominantly controlled by the circadian clock in light/dark cycles and that 48% were co-regulated by the circadian clock and light/dark cues (Figure 5a, Supplemental Data 1). Only 9% of the transcripts were primarily controlled by light/dark cues (Figure 5a, Supplemental Data 1). This is consistent with the expected underrepresentation of light/dark-controlled transcripts in a set of genes that oscillate in the absence of environmental cues.

**Figure 5:**
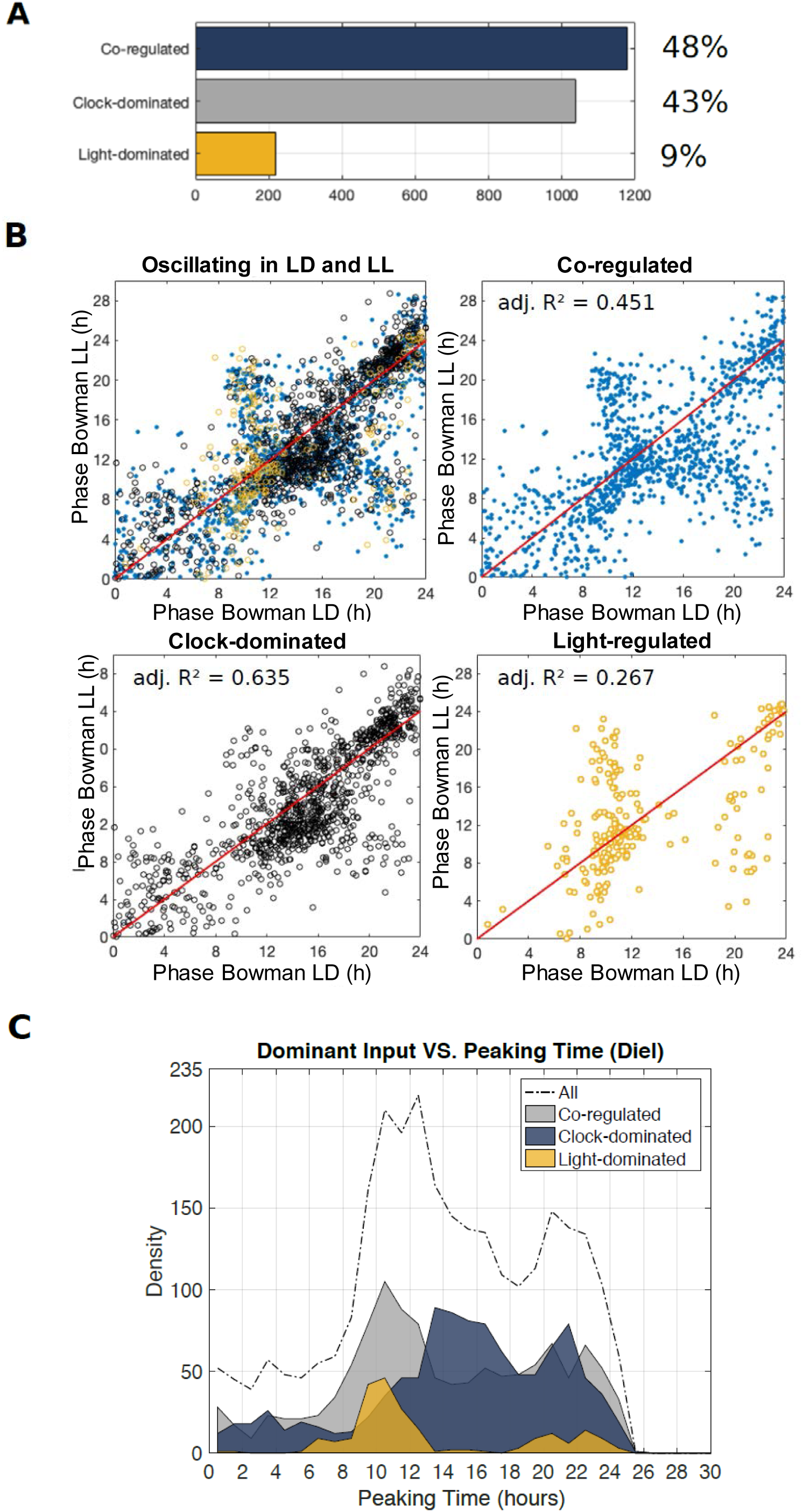
Relationship between external and internal cues to regulate the phase of the barley transcriptome **a)** Fractions of transcripts identified as clock-dominated, co-dominated by the clock and light and light-dominated by the Bode-analysis. **b)** Phase relationship between diel cycles (ND) and constant light (LL) for all transcripts oscillating in LD and LL and those dominated by the circadian clock, co-regulated by the circadian clock and light and light-dominated. **c)** Phase distribution of clock-dominated, co-regulated and light-dominated transcripts in diel cycles (ND).

We next investigated the phase relationship between driven and free-running conditions for transcripts predicted to be under the clock control, light control, and co-regulation by light and the clock by the Bode analysis (Figure 5b, c, Supplemental Data 1). The clock-dominated transcripts revealed the highest correlation (R^2^=0.64, Figure 5b) and the light dominated transcripts the lowest correlation (R^2^=0.27) of the phase between day/night cycles and constant light (Figure 5b). The correlation of the phase of co-regulated transcripts under LD and LL lay in between (R^2^=0.45, Figure 5b). This suggested that transcripts dominated by the circadian clock maintained a similar expression phase under changing light conditions, whereas the phase of transcripts dominated by light cues reflected the changes in light. These findings suggested that the Bode analysis predicted the main regulatory principles that determine the phase of oscillating transcription in day/night cycles. Namely, it suggested that about 40% of the transcripts with clock-maintained oscillations reveal a phase dominated by the circadian clock in diel cycles. For the remaining 60% of the transcripts with clock-maintained oscillations, the peak of their expression is under the control of light signaling pathways or co-regulated by light signaling and clock. This finding highlights the importance of light signaling pathways to regulate the phase of oscillating transcription even for the transcripts, the rhythmicity of which is maintained by the circadian clock.

## Discussion

The circadian clock was estimated to control a large proportion ∼25% of the of the transcriptome under constant conditions which is comparable to estimates for the proportion of circadian transcripts in Arabidopsis, rice (*Oryza sativa*) and poplar (*Populus trichocarpa*) (Filichkin et al. 2011, Covington et al. 2008, Michael et al. 2008, Gehan et al. 2015). Despite the strong control of the clock on transcript oscillations under LL, day-night cues had a major influence on shaping expression patterns of circadian transcripts under diel conditions. First, the expression phase under LL conditions was generally not a strong predictor of the transcript phase under LD conditions. The expression phase of transcripts was therefore a plastic trait where LD conditions delayed or advanced expression phase as compared to LL depending on the transcript. Second, the Bode analysis demonstrated that the majority of circadian transcripts was regulated by light/temperature or a combination of the clock and light/temperature cues under LD conditions. It is well known that the circadian clock is dynamically plastic and constantly entrained by metabolic and environmental cues for synchronisation with the cycles of the environment (Webb et al. 2019). Here, however, we demonstrate, that day-night cues do not only entrain the clock but can largely compensate for the lack of a functioning oscillator. The *hvelf3* and *hvlux1* mutants with no cycling transcriptome under LL conditions, were characterized by transcriptome oscillations under LD comparable to wild type Bowman. In this context it is interesting to note, that *hvelf3* and *hvlux1* mutants with a disrupted circadian clock, have been used to breed for barley cultivars adapted to Northern European environments with strong diurnal and seasonal changes in light and temperatures (Faure et al. 2012, Campoli et al. 2013, Pankin et al. 2014). Neither of the two arrhythmic mutants (*hvelf3*, *hvlux1*) have been reported to display any obvious impairment in photosynthesis and growth under conditions of pronounced photo- and thermocycles in contrast to the corresponding Arabidopsis mutants (Faure et al. 2012, Campoli et al. 2013, Habte et al. 2014). Similarly, Izawa et al. (2011) have reported that an *osgi* mutant in the field was not affected in photosynthesis and yield. Only under atypical growing conditions with late transplanting dates in the field, fertility was significantly reduced in *osgi* plants, indicating a loss of seasonal adaptability. Our data suggested that diel cycles could compensate circadian defects in the barley clock mutants, increase the number of oscillating transcripts compared to free-running conditions and strongly influence the time point of transcript peak expression. These findings suggested that the circadian oscillator has limited control over expression dynamics of circadian transcripts under conditions of pronounced diel oscillations in barley.

While the number of cycling transcripts was not different between the *hvelf3* and *hvlux1* mutants and Bowman, we observed quantitative variation in the phase distribution under diel conditions between the three genotypes. *HvELF3* altered the timing of transcript oscillations in day/night cycles by suppressing expression at the light and dark interfaces. This effect was apparently completely or partially independent of the oscillator defect that causes arrhythmia in the *hvelf3* plants under LL. Loss of *HvELF3*, but not of *HvLUX1*, altered the expression phase under diel cycles although both mutants had a disrupted circadian clock. Therefore, HvELF3 modifies the light and temperature controlled diel transcriptome oscillations in barley. The role of HvELF3 in mediating light and temperature cues is supported by the loss of photoperiod sensitivity in the *hvelf3* mutant (Faure et al. 2012) and resembles the role of ELF3 in antagonizing light input to the clock during the night in Arabidopsis (McWatters et al., 2000; Covington et al. 2001; Thines and Harmon, 2010). It is interesting that only HvELF3 but not HvLUX1 had strong effects on the diel transcriptome, because out of the Arabidopsis core components of the Evening Complex (EC) ELF3-ELF4-LUX only LUX has been identified as a transcription factor with direct DNA binding activity (Helfer et al. 2011). The different effects of *hvelf3* and *hvlux1* on the diel transcriptome may also be caused by the different nature of the underlying mutations, while the *hvelf3* mutant line carries a premature stop codon leading to a truncated HvELF3 protein, the *hvlux1* mutant is characterized by a single amino-acid exchange in the Myb-domain which is important for the binding to cognate DNA sequences and regulation of their target genes (Faure et al. 2012, Campoli et al. 2013). On the other hand, chromatin immune precipitation experiments demonstrated that ELF3 had many more significant binding sites than LUX suggesting that ELF3 also binds independently of LUX (Ezer et al. 2017). Therefore, HvELF3 and HvLUX1 might have independent targets in the barley genome.

Based on RNA time-series data we modeled a possible barley clock as a basis for understanding its effects on physiology, metabolism, and agronomic performance. It is important to emphasize that the resulting interactions between the individual components of the clock represent one of the possible solutions of the barley circadian clock circuit, which may serve as a null model in future studies aimed to experimentally resolve composition and regulation of this clock. We used simple dynamical models to capture gene regulatory dynamics without making *a priori* assumptions on the structure of the network. These dynamical models have been successfully used in the past to describe circadian processes of Arabidopsis under conditions that are similar to those of our dataset (Dalchau et al., 2012; Herrero et al., 2012; Banos et al., 2015; Mombaerts et al., 2016 and Mombaerts et al., 2019). It is also important to stress that our approach could only model genes with circadian expression oscillations, while it is well known that posttranscriptional regulation and the rate of protein degradation and activity is an essential constituent of the clock mechanism in Arabidopsis (Kim et al., 2003, Más et al., 2003).

Our modeling strategy used Hv*LUX1* to reveal the circadian circuitry, which therefore appeared as a major hub in the barley clock. Nevertheless, this predicted central role of Hv*LUX1* is consistent with the loss of self-sustained rhythms in the *hvlux1* mutant. Unlike Hv*ELF3* and Hv*ELF4,* Hv*LUX1* comprises known DNA binding domains suggesting that the transcriptional regulation of the EC converges on Hv*LUX1* (Nusinow *et al*., 2011). Our model predicted that Hv*LUX1* represses Hv*GI* and is itself repressed by HvLHY, consistent with the suggested repression of HvGI by the EC and *CCA1/LHY* repressing the Evening Complex in Arabidopsis (Fogelmark and Troein, 2014 Hsu *et al*., 2013).

Further, the regulatory predictions suggested that Hv*LHY* and Hv*RVE8* are activators of Hv*PRR73* and Hv*PRR95* in the morning and, at the same time, repress Hv*LUX1*. The morning activation of the Hv*PRRs* through *HvLHY* and Hv*RVE8*, together with the repression of Hv*LHY* and Hv*REV8* through the Hv*PRRs* later in the day, are also a key regulatory principle of the Arabidopsis clock (Hsu *et al*., 2013; Fogelmark and Troein, 2014). This suggests that the regulatory links between Hv*LHY*, Hv*RVE8*, and the Hv*PRRs* are conserved between barley and Arabidopsis, despite the independent evolutionary history of LHY-like and PPR-like genes in the barley and Arabidopsis clades (Takata *et al*., 2009, 2010, Campoli et al. 2012).

Our model suggested that Hv*PRR73*, the first *PRR* expressed in barley in the morning, activates Hv*PRR95*, which, in turn, activates Hv*PRR59* such that Hv*PRR73*, Hv*PRR95* and Hv*PRR59* are expressed in a sequential cascade. This resembles predictions by Pokhilko et al. (2011) who described the PRRs as a series of activators in the Arabidopsis clock, while other models have predicted that direct interactions between the PRRs are negative and directed from the later PRRs in the sequence to the earlier ones (Carré and Veflingstad 2013, Huang et al. 2012, Fogelmark and Troein 2014). However, the sequential regulation of the PRRs during the day appears to be a common feature of the circadian clock in both barley and Arabidopsis, while the sequence of expression of *PRR* genes is altered between Arabidopsis and barley. In Arabidopsis, the sequence of *PRR* expression starts with *PRR9* and ends with *PRR5*, while it the sequential *PRR* expression wave started with *PRR73* and ended with *PRR59* in our data (Hsu *et al*., 2013). Interestingly, *PRR37,* the major photoperiod response gene *PPD1* in wheat and barley, showed no circadian oscillations and is therefore probably not part of the circadian clock in barley. This is consistent with the finding that mutations in *PPD1* do not affect the circadian clock in barley and wheat (Campoli et al. 2012, Shaw et al. 2012).

Our modelling placed several members of the REVEILLE (RVE) gene family including barley homologs of the principal clock activators RVE8/6/4 into the barley clock (Kuno et al., 2003, Zhang et al., 2007, Rawat et al., 2009). The mutation in *eam7* had a limited effect on the central oscillator, but several RVE homologs lost rhythmicity or were strongly downregulated and this correlated with a lengthening of the period. The suggested that *eam7* is a component of the slave oscillator that only regulates a subset of clock-controlled transcripts. RVE-like genes have been implicated in such slave oscillators (Kuno et al. 2003) and *rve* mutants are characterised by a period lengthening (Hsu et al. 2013), however, none of the expressed *RVE* genes carried mutations that would alter the protein sequence.

BBX19, RVE7, and HAM3 were identified as three new candidate oscillator components in barley. While the three genes have already been proposed to have connections to the Arabidopsis oscillator, they have not been modeled as an integral part of the circadian clock but rather as clock outputs in Arabidopsis (Kuno *et al*., 2003; Wang *et al*., 2014). BBX19 acts as a gatekeeper of EC formation by mediating degradation of ELF3 and is part of a regulatory loop with CCA1 and/or LHY (Wang et al., 2015, Edwards et al. 2017, Tripathi et al. 2017) supporting the link between BBX19 and LHY in our model. The model also predicted that barley homologs of BT2, *CYP450*, *PHR1* are part of the core circadian oscillator in barley. However, these genes were only predicted to regulate other clock components and were not regulated themselves by clock genes. They therefore displayed a low connectivity within the circadian network, consistent with their known functions outside the central clock (Ren et al. 2007, Bak et al. 2011, Bari et al. 2006). Therefore, these components might provide input into the circadian network but are probably not components of the barley oscillator. Further work will be required to test the hypothesis on additional clock genes and interactions generated by the network modeling.

## Conclusion

Our comparison of diel and circadian transcriptomes in the different barley clock mutants revealed that fluctuations of light and temperature have a major effect on the diel oscillating transcriptome and can compensate for circadian defects in arrhythmic barley clock mutants. HvELF3, but not HvLUX1 controlled the expression phase of a large number of transcripts under diel conditions and this effect was independent from the oscillator arrest under LL. Dynamical modeling suggested novel putative clock genes and connections between clock genes as a basis for experimental explorations into the nature and functions of the barley circadian clock. Finally, our findings and the dataset provide a valuable resource for mining for the output targets of the barley clock genes HvELF3, HvLUX1 and EAM7 and to understand the role of the diel cues and clock in controlling the barley transcriptome and plant performance.

## Material and Methods

### Genetic material and growth conditions

Four spring barley genotypes were used in this study, the wild-type spring barley Bowman (BW WT) and three derived introgression lines with mutations in *HvELF3* (BW290), *HvLUX1* (BW284) and *EARLY MATURITY 7 (EAM7*) (BW287) (Faure *et al*., 2012; Campoli *et al*., 2013). BW290 carries an introgression of the *early maturity 8* allele (*eam8.k)* which is characterized by a base-pair mutation leading to a premature stop codon in *HvELF3*, which is orthologous to *ELF3* in Arabidopsis (Faure et al., 2012, Hicks et al. 2001). BW284 carries the *eam10* locus characterized by a single non-synonymous nucleotide polymorphism in the conserved Myb-domain of the barley LUX/ARRHYTHMO homolog (Campoli et al. 2013). The candidate gene for *EAM7* has not yet been identified but *eam7* is characterized by accelerated flowering and reduced sensitivity to the photoperiod (Stracke and Börner 1998). The lines were sown in soil (Einheitserde) in 96-well format. Seeds were maintained at 4°C for three days, followed by germination in 12 h light/12 h dark photoperiods at 20°C with a photon flux density 300µmol m^-2^s^-1^ during the day and 18°C during the night and grown for three weeks.

### Sampling, extraction of total RNA and sequencing

After three weeks of growth the second expanded leaf after the cotyledon leaf was harvested for two replicate plants per genotype and time point every 4h for 24h starting with lights off except for dusk and dawn when extra samples were taken for Bowman every two hours (Supplemental Figure 1). After completion of a 12h dark/12 h light diel cycle, the growth chamber was switched to constant light and 20°C and sampling continued for 36 h more starting from the first subjective dusk. Total RNA was extracted from ground tissue using a hybrid protocol of TRIZOL (Invitrogen) and purification columns from a RNeasy RNA extraction kit (Qiagen). Extracted total RNA was DNAse treated (Ambion). The concentration and integrity of the extracted RNA was determined on a BioAnalyzer (Agilent) before library preparation. The library preparation was carried out following the TruSeq protocol and single-end sequenced on a HiSeq 2500 with 10 Million reads per library.

### Mapping of reads and rhythmicity analysis

The quality of the sequencing data was verified using the FastQC software. The reads were mappedagainst a custom barley reference transcriptome (Digel et al., 2015) and raw read counts were obtained using the software implementing the full pipeline for RNA-seq analysis RobiNA (version 1.2.3) with default settings (Lohse et al., 2012). Raw read counts were normalized to counts per million (CPM) using the R/Bioconductor package “edgeR” and used for the downstream rhythmic analysis and modeling. To cross-reference a nomenclature of the gene models used in this study (Digel et al., 2015) with the identifiers of the HORVU gene models annotated on the barley pseudochromosomes (Mascher et al., 2017), we used reciprocal blastn v. 2.9.0+ (e-value cut-off 10e-05). The reciprocal best blast hit pairs were extracted as matching gene model identifiers.

For the analysis of the day/night data, the sequence of samples was inverted to start with the night followed by the day samples (12h, 14h, 16h, 20h, 0h, 2h, 4h, 8h), and this data set was therefore termed night-day (ND). The oscillating patterns of gene expression and period (as a duration of one complete cycle) and phase (as the location of time of the peak of the curve) of the curves were determined using the ARSER algorithm in the R package “MetaCycle” (Wu et al., 2016; Yang et al., 2010). To increase the analytical power for the rhythmic analysis, the 24 h diel dataset, but not the 36 h LL data, was duplicated to imitate 48 h of sampling data. The settings for the ARSER algorithm were adjusted for each genotype individually so that the period was normally distributed with two symmetric tails and the number of transcripts passing the cut-off (Benjamini-Hochberg corrected false discovery rate of 0.1) was maximal with the given range of period estimation being minimal. For Bowman and BW287 the settings are shown in Figures 2b and 2c and comprised mean = 28, upper/lower limit =22/34 for the constant light data. In diel cycles, the mean and upper/lower limits were set to 24 and 21/27, respectively.

### Modeling the barley circadian clock

In order to identify putative barley clock genes, we blasted all expressed barley transcripts against the Arabidopsis transcriptome (geneontology.org). For each barley transcripts the best Arabidopsis blast hit was retained and we then selected only those barley transcripts for which the Arabidopsis homolog carried the GO annotation “circadian” and “transcription” (Supplemental Data 1). The resulting 138 barley transcripts were then further filtered for those that exhibited unambiguous dynamics and a high signal-to-noise ratio of expression. This filtering step was necessary to ensure that we did not identify dynamics out of noise. Hence, genes for which the amplitude of oscillation were lower than 20 CPM in the last 24 hours were removed. The choice of such filter is motivated by both the transitional nature of constant light data, which typically shows a large decrease of amplitude after few hours in barley, and the dependency of noise on gene expression levels. Furthermore, genes that were constantly up/down regulated without exhibiting further significant dynamics were also discarded. This was performed by detrending the 24 last hours of constant light data before applying the same filtering criterion. After filtering, out of 138, 49 and 48 genes passed the filtering criterions in WT and BW287 datasets, respectively. BW284 (*Hvlux1*) and BW290 (*Hvelf3*) datasets were not considered in the following network analysis since these mutations led to the arrhythmic transcriptomes. Finally, seven genes (*Hv.21080, Hv.22191, Hv.23289, Hv.32914, Hv.33010, Hv.6793, MLOC_7084.3*) were manually discarded from both subsets list of candidates as they were not DNA binding transcription factors but rather enzymes in a metabolic process, leaving 42 in Bowman and 41 in BW287 of which 35 transcripts were in common between the Bowman and BW287 final datasets used for modeling. The HvELF3 transcript did not pass the filtering and, therefore, could not be used to infer dynamical interactions.

To model the barley circadian clock based on the time-course gene transcription data, we adopted an approach based on Linear Time Invariant (LTI) models. LTI models do not rely on prior knowledge of the transcriptional network to provide accurate predictions and havebeen shown to provide reliable predictions of the dynamical processes involved in the Arabidopsis circadian network (Dalchau et al., 2012; Herrero et al., 2012; Mombaerts et al., 2016 and Mombaerts et al., 2019). To provide a comprehensive evaluation of the LTI model, the performance of the modeling strategy was evaluated and compared under conditions that replicated those of the experiments using widely used benchmarks models (Supplemental Information).

We used first order models to represent the system dynamics between two genes at a time using an LTI model with the following equation:

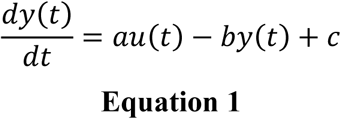

With *by*(*t*) corresponding to the degradation rate of y and *au*(*t*)that represents the influence of another transcription factor through the synthesis rate of y. The model, therefore, evaluates whether the rate of change of a particular gene *y* depends on another gene *u*. Estimating a model means finding (a), (b) and (c) that produce a vector y(t) as close as possible to the real data. The estimation of parameters was performed using the function ‘pem’ implemented in MATLAB that minimizes the prediction error of the data. LL data were used for the estimation, as they represent the autonomous behavior of the oscillator.

The goodness of fit of the model with the data was calculated as following:

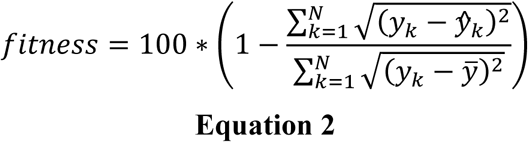

Where *y_k_* is the data (output), *ȳ* is the average value of the data, and *ŷ_k_* is the estimated output. MATLAB function *compare* was used to compute the fitness of the model. Each potential link between two genes was validated if the associated model reproduced the dynamics involved with a sufficient degree of precision, which corresponds to a fitness threshold estimated at 60% (Supplemental Information).

To investigate the potential regulators of *HvLUX1*, a collection of independent 1^st^ order LTI models was estimated separately between each of the transcript and *HvLUX1* in the Bowman background. In each case, the parameters were estimated so that they together provide the best possible fit to the HvLUX1 time-course data. This step takes the following form:

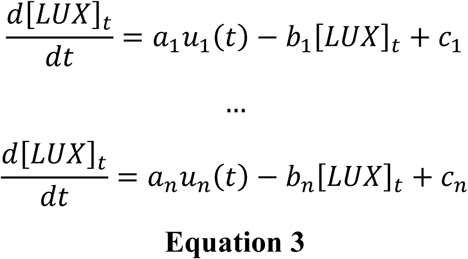

Where *n* corresponds to the number of candidates (42 models in total). Each model was characterized by a fitness metric that ranges from 0 to 100%, representing its capability to describe the regulatory dynamics between genes. A gene, therefore, would be further considered as a regulator for HvLUX1 if the model is capable of reproducing the shape of *HvLUX1* with a sufficient degree of precision. A fitness threshold, evaluated from *in-silico* benchmarks systems (Supplemental Information), was used to validate the models. In this case, the fitness threshold was set to 60% to limit false positive predictions of regulatory interactions while accounting for sufficient gene regulatory models to describe the system of interest. Finally, 20 models passed the validation step (Supplemental Data 6). The methodology is summarized in Supplemental Figure S4.

To further narrow down the predicted regulatory interactions, we estimated the consistency of the candidate models using the filtered *eam7* (BW287) dataset. For this purpose, we evaluated 1^st^ order LTI models for each of the previously identified regulations and retained those with the goodness of fit > 60% in the *eam7* (BW287) experimental condition (Supplemental Figure S4). To keep links with the highest confidence only, the dynamical consistency of the LTI models based on these two independent datasets (Bowman and BW287) was evaluated using the nu (ν) gap metric (*gapmetric,* MATLAB) (Vinnicombe, 1993; Mombaerts *et al*. 2019). We further considered models that had a v-gap less than 0.2 following Carignano et al. (2015). As a result, six regulatory interactions were filtered out (Hv.10528 to Hv.27754, Hv.1530 (GI) to Hv.19411, Hv.19411 to Hv.20312 (LUX), Hv.19759 (TOC1) to Hv.20312 (LUX), Hv.5253 (LHY) to Hv.27754, Hv.9855 to Hv.18813 (PRR59)). HvPRR95 (Hv.4918) appeared as a hub with 8 connections so that we repeated the search for regulators of HvPRR95 (Equation 3), computed their interactions in both datasets, and checked their consistency.

The relative contribution of light signaling and circadian clock pathways in generating oscillating transcriptome was evaluated using a Bode analysis (*bode* function in MATLAB) with the threshold of 7 dB to discern between the two alternative regulatory inputs (Dalchau *et al*., 2010) (Supplemental Figure S5). Here, we use the magnitude response of the signal to assess the relative contribution of the inputs *u_light_*(*t*) and *u_LHY_*(*t*) in each of the validated model, at a frequency of 24h (or .262 rad h ^-1). Following Dalchau et al. (2010) if the magnitude of the response of the light input was 7dB higher than the contribution of the clock (represented by *HvLHY* potentially delayed), the circadian regulated gene (the output of the model) was considered driven mostly by light; in the opposite case, the transcript expression was considered as driven by the clock. If the magnitude difference was less than 7dB, then the circadian regulated gene was considered regulated by both inputs equally. The methodology is summarized in Supplemental Figure S7B.

To this end, we used the 2759 transcripts that were identified as oscillating in both diel and free-running conditions in the wild-type Bowman background to calculate another set of LTI models as described earlier. As a reference, we selected a formerly identified clock gene peaking in the morning, *HvLHY* (Hv.5253), with a range of delays integrated into the model to implicitly represent the clock input following Dalchau et al. (2010). The structure of such models is schematically represented in Supplemental Figure S7A. This way, the light input is incorporated on two levels: explicitly through the light input and implicitly through the clock pathway using the following equation:

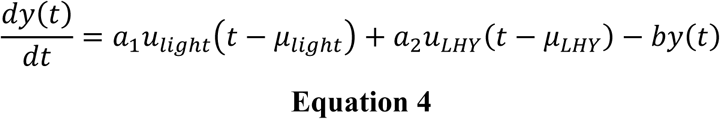

Where *u_light_* was assumed to be binary (1 = light; 0 = dark). We fixed the light delay *µ_light_* to 0h to represent the effect of rapid light signaling on the transcripts, and computed delays ranging from 0 to 8h, every 0.2h, for *HvLHY*. The delay that provided the best fit to the data was selected independently for each transcript. Ultimately, models were validated if they succeeded in capturing the regulatory dynamics involved with a goodness of fit > 60%.

We assessed the accuracy of our LTI-based network reconstruction algorithm on the circadian model from Pokhilko et al. (2010) as detailed in the Supplemental Information. The performance of the modeling strategy was evaluated based on the Area Under the curve of the Receiver Operating Characteristic (AUROC) and Area under the Precision-Recall Curve (AUPREC) criteria (Supplemental Information, Supplemental Figure S8).

The scripts for the modelling of the barley clock are available under: https://github.com/Lmombaerts/CircadianBarley

## Accession numbers

ArrayExpress accession E-MTAB-8372

## Acknowledgements

We cordially thank Kerstin Luxa, Caren Dawidson, Thea Rütjes and Andrea Lossow for excellent technical assistance.

## Authors’ Contribution

M.K. and S.J.D. conceived the original research project. L.M.M. and M.K. designed the experiments. L.M.M. carried out the experiments and analysed the RNA-sequencing data. L.M. calculated LTI models with the help of J.G. and A.W. A.P. contributed to the bioinformatic analyses. L.M.M., L.M., A.W., A.P. and M.K. wrote the manuscript.

